## Supplementary figures and images for "Real-time structural motif searching in proteins using an inverted index strategy"

### Supplemental Figure 1

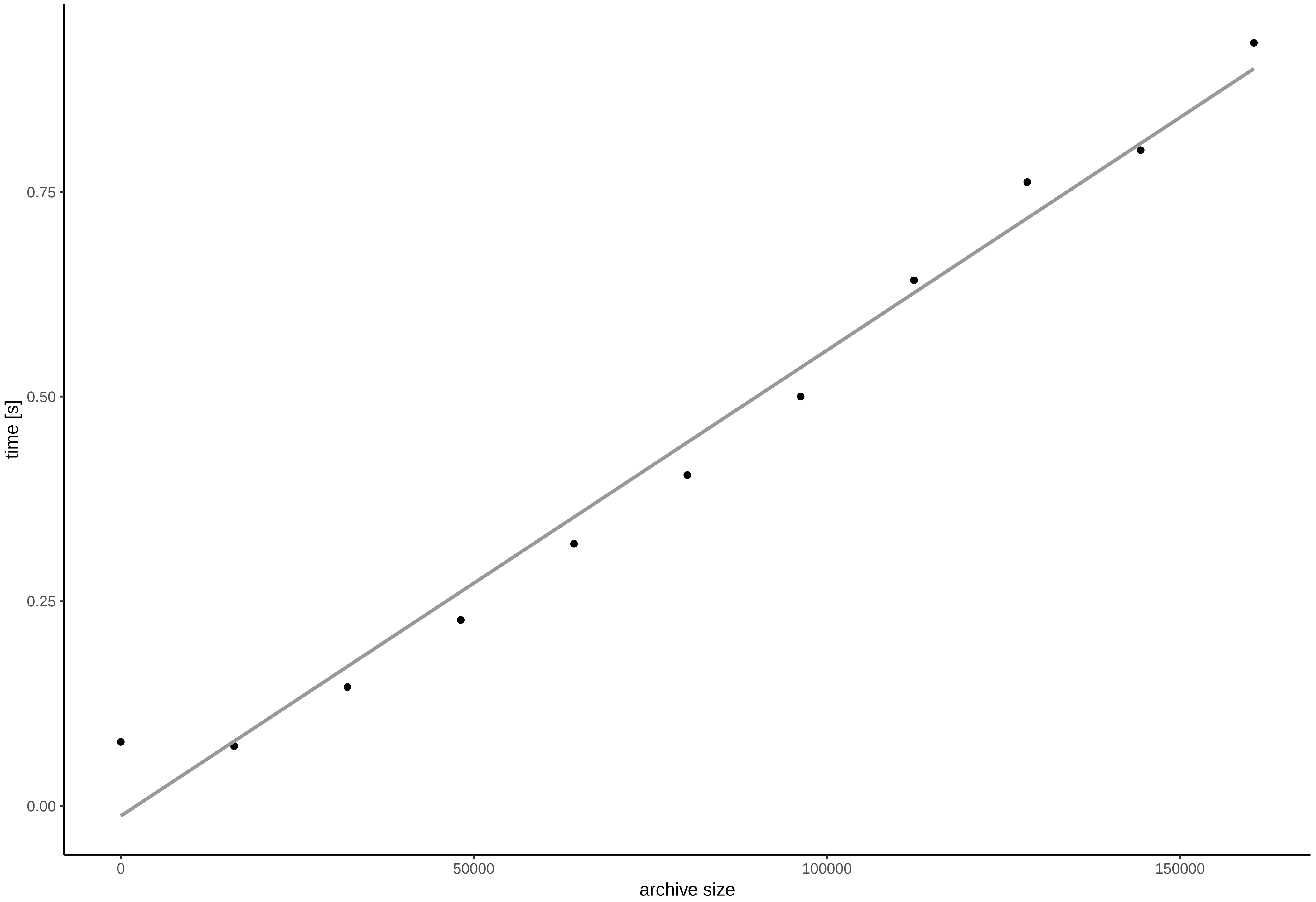

### Supplemental Figure 2

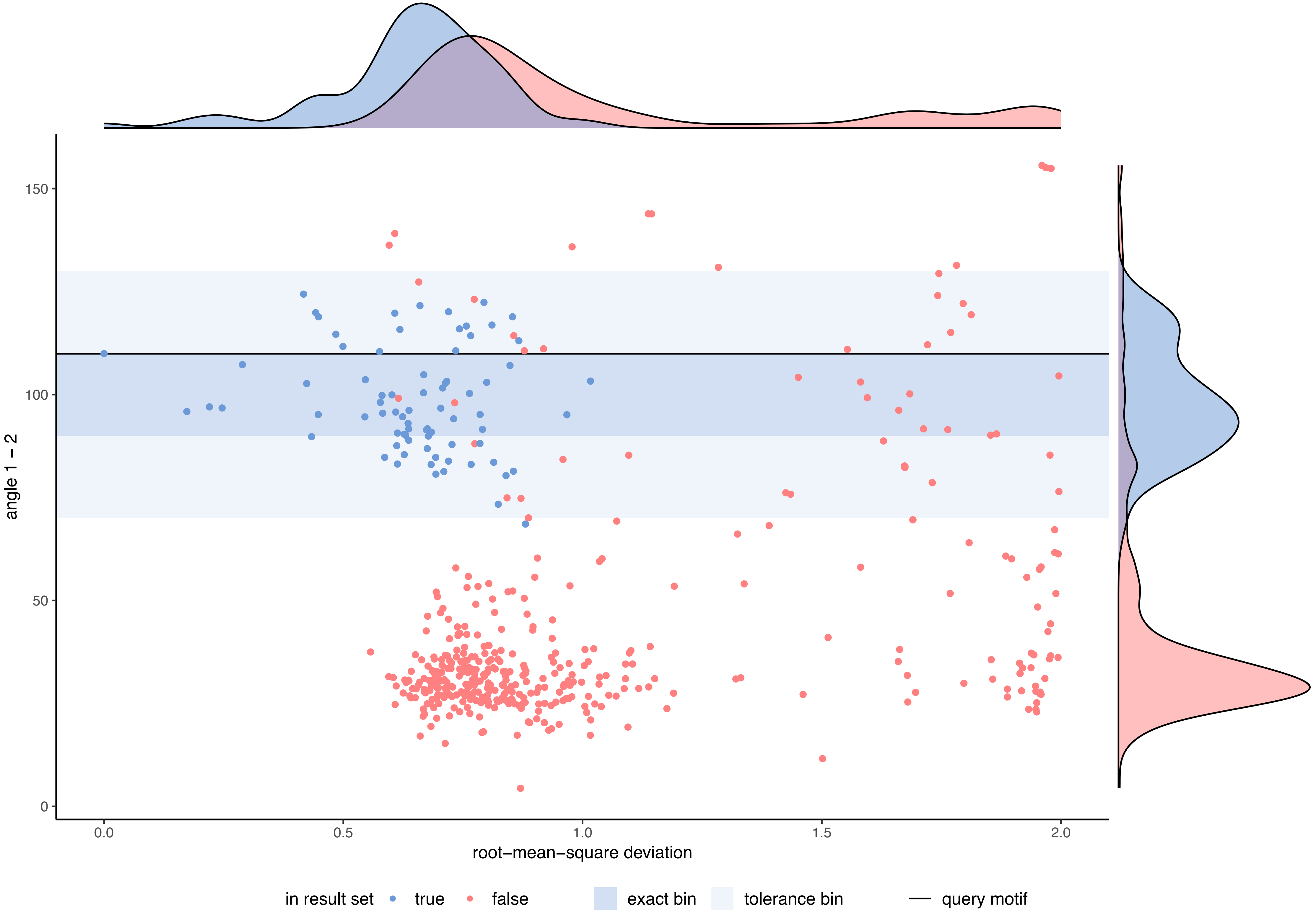

### Supplemental Figure 3

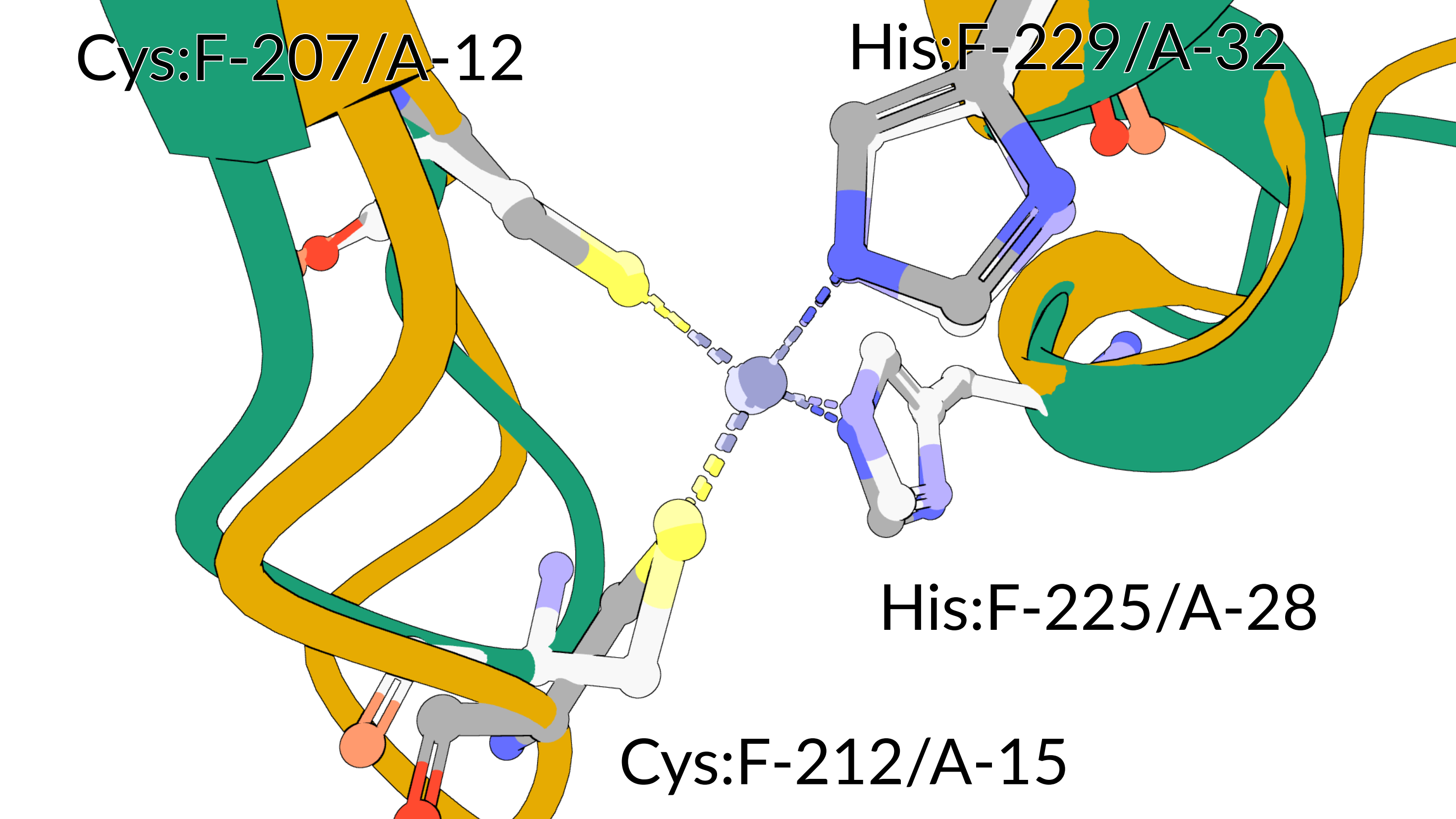
